# *Candida auris* is rendered non-viable by medium-chain fatty acids

**DOI:** 10.1101/2021.07.13.452291

**Authors:** Kalynne R. Green, Magdia De Jesus, Kearney T. W. Gunsalus

## Abstract

The medium-chain fatty acids, octanoic and decanoic acid, found in coconut oil, were fungistatic and decanoic acid was fungicidal against a panel of *Candida auris* strains, during both planktonic and biofilm growth. The strains were from all four major geographic clades, and some were resistant to several classes of antifungal drugs. These compounds are safe, natural products and could provide a new strategy for skin decolonization and environmental decontamination.

---

*Candida auris* has rapidly emerged around the world as a drug-resistant fungal pathogen causing hospital outbreaks with high mortality rates. Factors contributing to its virulence likely include *C. auris*’ persistence both as a skin colonizer and in the environment (1, 2); its ability to transfer to other patients or healthcare workers; and its resistance to various disinfection measures and antifungal drugs [reviewed in (3, 4)].

While the drug susceptibility profile varies between strains, and particularly between clades, many strains of *C. auris* are resistant to at least two of the three major classes of antifungal drugs, and the emergence of pan-resistant strains occurs during treatment with antifungals [reviewed in(3, 5)].

There is an urgent need for new methods to achieve environmental decontamination and particularly skin decolonization, without using antifungal drugs. Coconut oil, its derivative medium-chain triglyceride (MCT) oil, and their constituent medium-chain fatty acids (MCFAs), have antimicrobial properties against other *Candida* species *in vitro* (6–9). We showed that dietary coconut oil reduces gastrointestinal colonization by *Candida albicans* in mice (10), and that dietary supplementation with MCT oil reduces *Candida* GI colonization in humans (11). Coconut oil is safe for human consumption and topical application; it is widely used in cooking and as a skin and hair care product. In this study, we investigated the antifungal activities of the medium-chain fatty acids, octanoic and decanoic acid, against *Candida auris*.

## *C. auris* strains are susceptible to growth inhibition by medium-chain fatty acids

The antifungal activity of octanoic and decanoic acid was analyzed against a panel of strains of *C. auris* obtained from the Centers for Disease Control, specifically to be used in the laboratory of M. De Jesus (**Table 1**). The panel includes clinical isolates from four geographic clades (S. Asia, E. Asia, S. Africa, and S. America), both aggregate-forming and non-aggregate forming strains, and strains with resistance to azoles and amphotericin B. To work with the strains of *C. auris*, precautions were taken as previously described (2). The susceptibility of the *C. auris* strains to octanoic and decanoic acids was determined using a broth microdilution assay [(12) and supplemental methods]. Briefly, strains were seeded into 96-well plates in RPMI 1640 2% glucose in the presence of a two-fold dilution series of octanoic or decanoic acid (final concentration 0.02-10 mM) and grown for 24 hours at 37 °C. Sealed plates were read with a microplate reader (530 nm), and results were expressed as a percentage of growth compared to the untreated cells.

**Table 1.** Strains used in this study.

| Strain no. | AR Bank # | Geographic clade<sup>a</sup> | Aggregation<sup>b</sup> | MIC (mg/L)<sup>a</sup> |  |  |  |
| --- | --- | --- | --- | --- | --- | --- | --- |
|  |  |  |  | Amphotericin B (≥2 mg/L) | Caspofungin (≥2 mg/L) | Fluconazole (≥32 mg/L) | Flucytosine |
| CAU-01 | 0381 | II (East Asia) | No | 0.38 | 0.125 | 4 | 2 |
| CAU-02 | 0382 | I (South Asia) | Yes | 0.38 | 0.5 | 16 | 0.125 |
| CAU-03 | 0383 | III (South Africa) | Yes | 0.38 | 0.25 | 128 | 0.5 |
| CAU-04 | 0384 | III (South Africa) | Yes | 0.5 | 16 | 128 | 0.5 |
| CAU-05 | 0385 | IV (South America) | Yes | 0.5 | 0.5 | >256 | 0.5 |
| CAU-06 | 0386 | IV (South America) | Yes | 0.5 | 0.5 | >256 | 0.5 |
| CAU-07 | 0387 | I (South Asia) | No | 0.75 | 0.25 | 8 | 8 |
| CAU-08 | 0388 | I (South Asia) | No | 1.5 | 1 | >256 | 0.125 |
| CAU-09 | 0389 | I (South Asia) | No | 4 | 0.5 | 256 | 128 |

<sup>a</sup> From the CDC & FDA Antibiotic Resistance (AR) Isolate Bank *Candida auris* (CAU) panel (https://www.cdc.gov/ARIsolateBank/Panel/PanelDetail?ID=2). MICs obtained by broth microdilution. CDC's tentative breakpoints for each drug are provided (CDC. *Candida auris*: information for laboratorians and health professionals. Atlanta, Georgia: US Department of Health and Human Services, CDC; 2019. https://www.cdc.gov/fungal/candida-auris/health-professionals.html), and resistance highlighted.
<sup>b</sup> (19)

Regardless of geographic clade or antifungal susceptibility profile, growth of all strains was inhibited by MCFAs. During planktonic growth, octanoic acid elicited 80% growth inhibition (pMIC80) at 2.5-5 mM, and pMIC50 was 0.6-2.5 mM, depending upon the strain (**Figure 1A**). Decanoic acid is more potent, with a pMIC80 of 1.25-2.5 mM, and a pMIC50 of 0.6-1.25 mM (**Fig 1B**).

**Figure 1.**
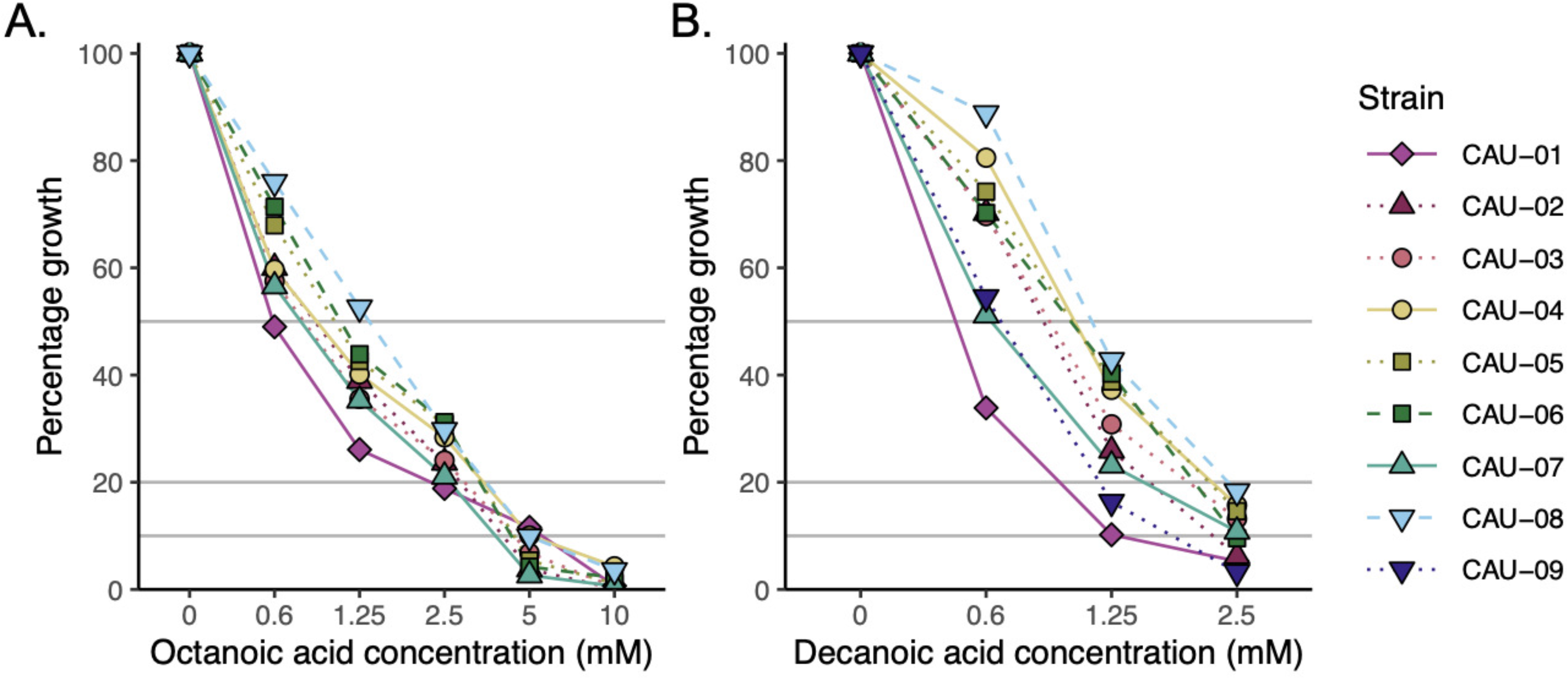
*C. auris* strains are susceptible to growth inhibition by octanoic and decanoic acids. Strains were grown in the presence of indicated concentrations of **A)** octanoic or **B)** decanoic acid for 24 hours.

## Antifungal-resistant *C. auris* strains are rendered non-viable by decanoic acid

To determine whether octanoic and decanoic acids are fungicidal, or merely fungistatic, cells from the wells treated with concentrations of octanoic and decanoic acids at or above the pMIC50 were serially diluted tenfold and spotted onto Sabouraud dextrose agar to determine whether viable cells could be recovered. While none of the tested concentrations of octanoic acid resulted in complete loss of viability, recovered colonies displayed variable colony morphology (particularly decreased colony size in some strains, **Fig 2A**). In contrast, decanoic acid was fungicidal, with minimum fungicidal concentrations (the lowest concentration showing no growth) of 5 or 10 mM for all strains tested (**Fig 2B-C**).

**Figure 2.**
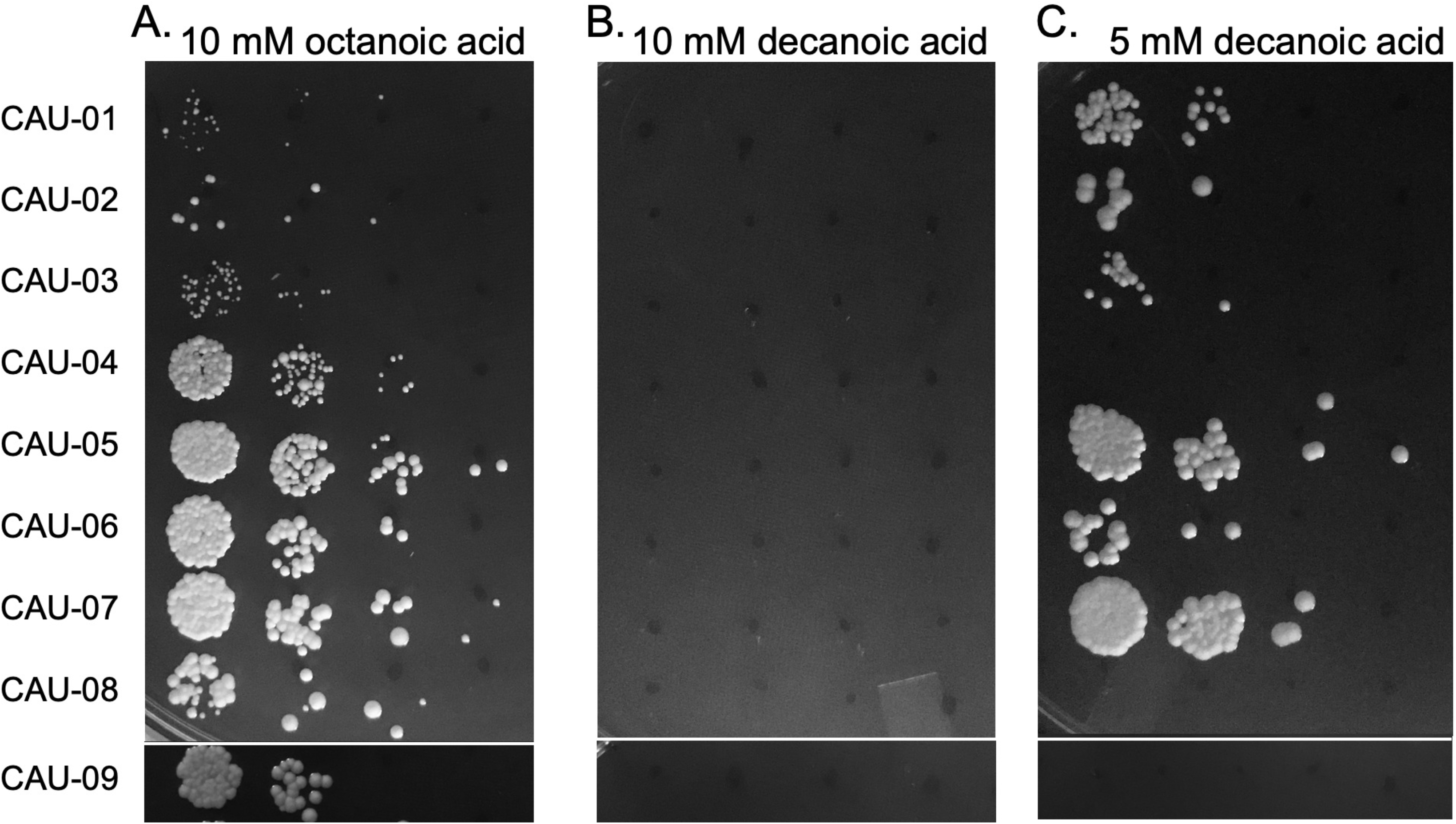
Minimum fungicidal concentration of decanoic and octanoic acids. Strains grown in the presence of **A)** 10 mM octanoic acid, **B)** 10 mM decanoic acid and **C)** 5 mM decanoic acid were serially diluted ten-fold, spotted onto Sabaraud dextrose agar, and incubated at 37 °C for 48 hours. (No additional colonies were observed after one week at 37 °C.)

## Octanoic and decanoic acids reduce the viability of *C. auris* biofilms

*Candida spp.* biofilms tend to be highly resistant to antifungal drugs, even for strains that are drug-susceptible during planktonic growth. To determine the susceptibility of mature *C. auris* biofilms to octanoic and decanoic acids, *C. auris* biofilms were grown for 24 hours and then treated with octanoic or decanoic acid for an additional 24 hours. Cell viability was measured by metabolic activity of the cells using an XTT reduction assay (adapted from (13); supplemental methods). The sessile MIC50 (sMIC50) for octanoic acid was 2.5-20 mM for eight of the nine strains tested. All strains showed a sMIC50 of 1.25-5 mM, and sMIC80 of 5-10 mM, for decanoic acid (**Fig 3**).

**Figure 3.**
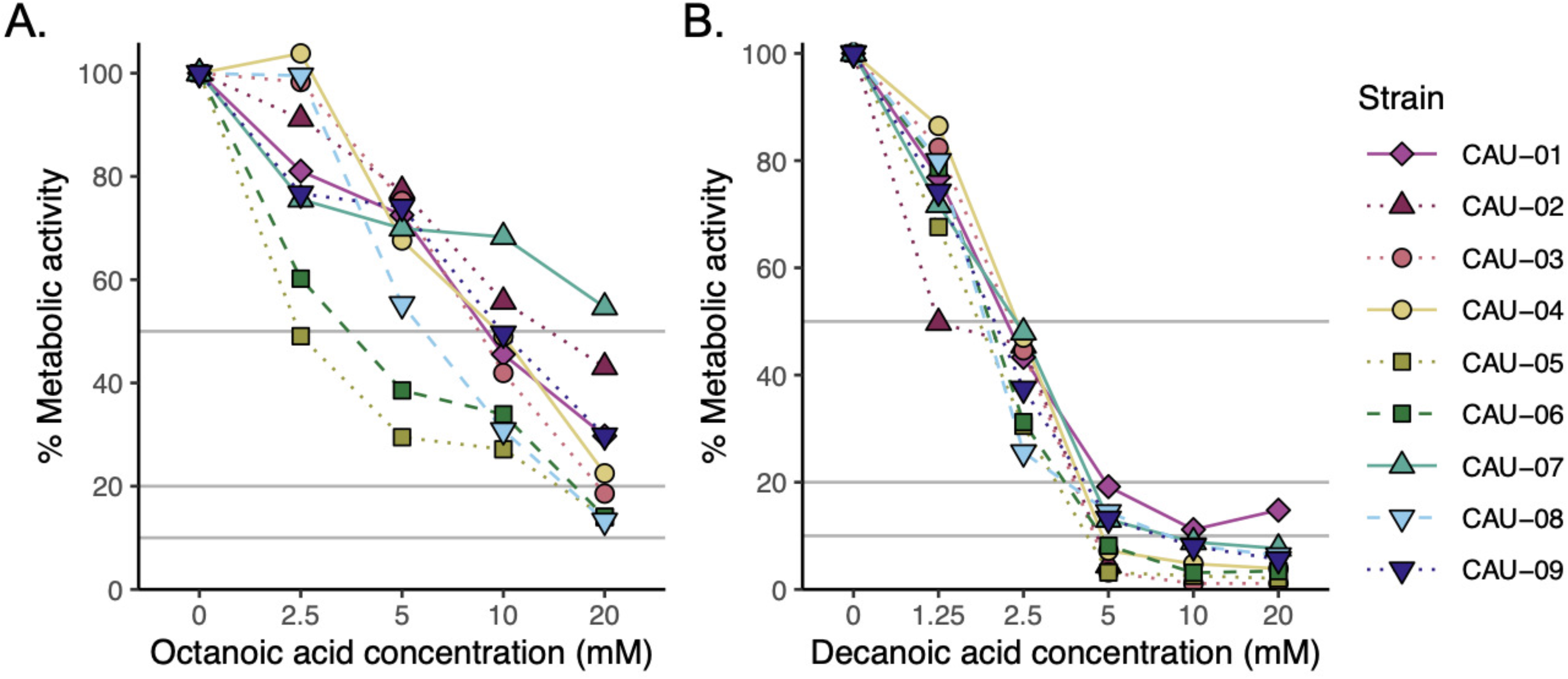
Susceptibility of *C. auris* biofilms to octanoic and decanoic acids. Biofilms were grown for 24 hours, exposed to indicated concentrations of **A)** octanoic or **B)** decanoic acids for an additional 24 hours, and cell viability measured via XTT reduction.

Medium-chain fatty acids exhibited antifungal activity against *C. auris* isolates from all four major geographic clades; strains resistant to various classes of antifungal drugs, and both aggregate-forming and nonaggregating isolates. Non-aggregate-forming isolates have greater pathogenicity (14). Regardless of clade or antifungal resistance, all strains were susceptible to growth inhibition and were rendered non-viable by medium-chain fatty acids; moreover, the medium-chain fatty acids can greatly reduce the viability of *C. auris* biofilms. Biofilm formation tends to confer additional resistance to antifungals (15, 16), including via the upregulation of drug efflux pumps (17) and the sequestration of antifungals in the extracellular matrix (18). This broad susceptibility to MCFAs suggests that resistance to currently-available antifungal drugs does not confer resistance to MCFAs, and that the mechanisms of antifungal activity are different.

There is an urgent need for safe ways to reduce skin colonization by *C. auris* without using antifungals. This research suggests a new strategy to address this rising threat. MCFAs have been shown to have antimicrobial properties against a range of microbes, including other *Candida* species. The MCFAs found in coconut oil are safe for human consumption and topical application, even young and ill individuals. MCT oil, refined from coconut oil, is rich in MCFAs including octanoic and decanoic acid, is commercially available as a nutritional food, and already has multiple medical applications. The results of this study demonstrate that derivatives of coconut oil that are widely available and known to be safe, can greatly reduce the viability of a broad spectrum of *C. auris* strains, including strains from all four major geographic clades, and during both planktonic and biofilm growth. The efficacy of the MCFAs against a diverse panel of *C. auris* isolates argues for the exploration of MCFAs as a more general antifungal strategy — one that may prove particularly valuable given the global emergence of drug-resistant fungi such as *C. auris*.

## Materials and Methods

### Planktonic susceptibility assay

Strains were pre-cultured onto Sabouraud dextrose agar. Octanoic and decanoic acids were obtained from Sigma Aldrich (O3907 and C1875) and stock solutions (2 M in 100% ethanol) were stored in glass vials at −80 °C. A two-fold dilution series (2-0.004 M; final concentrations 10-0.02 mM) of fatty acid was prepared in 100% ethanol, then diluted 1000x into 2x RPMI 1640 2% G (2% glucose; with L-glutamine and phenol red, without sodium bicarbonate; buffered to pH 7.0 with 165 mM MOPS [3-(N-morpholino) propanesulfonic acid]), which was pre-warmed to increase solubility of fatty acids. 100 μL was dispensed into each well of a flat-bottomed 96-well plate (CellTreat 229596); untreated wells contained an equivalent volume of ethanol (vehicle control). *C. auris* inocula were prepared by suspending a colony in sterile water and adjusting the OD_530_ to 0.011 (~1-5 × 10^5^ CFU/mL); 100 μL of inoculum was added to each well except the blanks (final inoculum ~0.5-2.5 × 10^5^ CFU/mL). Plates were incubated in ambient air at 37 °C for 24 +/− 2h, and OD_530_ was measured using a microplate reader (VersaMax, Molecular Devices). Data analysis was performed using R (20) and the *tidyverse* package (21).

### Biofilm susceptibility assay

Liquid cultures in Sabouraud dextrose broth were grown overnight at 37 °C, 150 rpm. Cells were pelleted via centrifugation, washed twice with and then resuspended in phosphate buffered saline (PBS, pH 7.4). The OD_530_ was adjusted to 0.11 (~1-5 × 10^6^ CFU/mL) in RPMI 1640 2% G, and 100 μL of inoculum was added to each well of a flat-bottom tissue culture-treated 96-well plate (Corning 3596), and incubated in ambient air at 37 °C for 24 hours. Medium was removed and wells washed with PBS to remove non-adherent cells. Biofilms were treated with a two-fold dilution series of octanoic or decanoic acid in RPMI 1640 2% G for 24 hours at 37 °C. Biofilms were washed with PBS; XTT solution (0.5 g/L, 1 μM menadione, in PBS) was added, plates were incubated for 2-3 hours at 37 °C in the dark, and absorbance at 490 nm was measured using a VersaMax microplate reader. The results for each strain were expressed as a percentage of the 0 mM fatty acid control. Data analysis was performed using R and *tidyverse*.

## Acknowledgments

We thank members of the De Jesus lab Steven Torres and Adina Longyear for their technical assistance, and Dr. Navjot Singh of the Wadsworth Center Applied Genomic Technologies Core for the helpful discussions of our data. We also thank Richard Cole of the Wadsworth Center Advanced Light Microscopy and Image Analysis Core. M. De Jesus laboratory was supported by the University at Albany and Wadsworth Center start-up funds. KRG was supported by the Summer Scholars program at Siena College. KTWG was supported by a Summer Faculty Research Fellowship from the Siena College Committee on Teaching and Faculty Development. The funders had no role in study design, data collection and interpretation, or the decision to submit the work for publication.

